# Deep-brain thermo-endomicroscopy

**DOI:** 10.1101/2025.11.25.690615

**Authors:** Ryuki Imamura, Yurina Nakane, Naoya Kataoka, Hiroshi Abe, Takeshi Ohshima, Shingo Sotoma, Takuma Sugi

**Affiliations:** Program of Biomedical Science, Graduate School of Integrated Sciences for Life, Hiroshima University, Higashi-Hiroshima, Hiroshima 739-0046, Japan; Department of Integrative Physiology, Nagoya University Graduate School of Medicine, Nagoya 466-8550, Japan; National Institutes for Quantum Science and Technology (QST), Quantum Materials and Application Research Center (QUARC), 1233 Watanuki, Takasaki, Gunma, 370-1292, Japan; Department of Materials Science, Tohoku University, Sendai, Miyagi, 980-8579, Japan; Graduate School of Science and Technology, Kyoto Institute of Technology, Hashikami-cho, Matsugasaki, Sakyo, Kyoto, 606-8585, Japan

## Abstract

Subtle deviations in biophysical parameters, particularly temperature, within deep-brain regions exert substantial effects on whole-body homeostasis and metabolism. However, the underlying mechanisms remain poorly understood because cellular-resolution measurements of these parameters in the deep brain have been technically inaccessible. Here, we present a lensless fiber-optic quantum-sensing endomicroscopy that enables cellular-resolution temperature mapping in the mouse deep brain. A microwire-coupled, lensless fiber bundle permits simultaneous temperature readout from optically detected magnetic resonance spectra of >50 fluorescent nanodiamonds, achieving cellular resolution (4.88 μm) with a precision of 0.49 ℃ (minimum precision, 0.18 ℃) within a 0.28 mm^2^ field of view. We demonstrated temperature mapping of multiple brain depths in head-fixed and vigorously behaving mice subjected to nociceptive stimulation, detecting lower temperatures during anesthesia compared to the awake state in ventromedial hypothalamic nucleus (VMH) at a 5 mm depth. These mappings reveal that spatio-temporal temperature distribution and its anesthesia-dependent reduction are heterogeneous rather than uniform in VMH. This endomicroscopy provides a promising platform to dissect how deviations in biophysical parameters modulate deep-brain function.

## Introduction

Deep-brain regions orchestrate whole-body homeostasis and metabolism. For example, hypothalamic nuclei coordinate thermoregulation, energy balance and arousal, whereas the suprachiasmatic nucleus functions as the master circadian clock, aligning daily rhythms of behavior and metabolism^1–3^. Subtle shifts in deep-brain biophysical parameters can modulate these regulatory processes. In particular, temperature is an internally and externally modulated parameter whose subtle deviations have tremendous influences on cellular ion-channel, receptor kinetics, synaptic transmission and network excitability^4–7^. The impact of deep-brain temperature is exemplified by studies demonstrating that manipulating hypothalamic temperature alters mouse lifespan^8,9^. Beyond temperature, acidosis within the amygdala represents another biophysical parameter capable of eliciting fear behaviors^10,11^. Nevertheless, despite the recognized importance of such parameters, their underlying mechanisms remain elusive because cellular-resolution measurements in the deep brain have largely been out inaccessible.

Technologies to measure biophysical parameters have been demonstrated in cultured cells, *Caenorhabditis elegans* and mice, but commonly used tools average away mesoscale structure in deep brain regions. Infrared thermography is restricted to surfaces and cannot provide cellular resolution readouts from subcortical targets^12^. Fiber photometry collapses fluorescence over a scattering-defined volume adjacent to the fiber tip even when using tapered or engineered fibers designed to segment collection along the shank, thereby precluding cellular-resolution temperature mapping^13–16^. MR thermometry and spectroscopy provide valuable organ- or region-level estimates but are constrained by calibration variability, voxel averaging, and restricted temporal and spatial resolution *in vivo*, preventing cellular mapping in the deep brain^17^. Conventional two-photon microscopy achieves cellular resolution but is fundamentally depth-limited in brain tissue (typically ≤ ∼1–1.6 mm in mouse cortex), leaving many deep-brain regions inaccessible^18^. Gradient index (GRIN) lens-based endomicroscopy is often used to image the deep brain, but its spatial resolution is typically compromised by the off-axis and depth-dependent aberrations characteristic of GRIN lenses^19^. At the probe level, protein-based fluorescent sensors suffer from photobleaching and cross-talk between multiple parameters, such as pH and ion concentration.

Quantum sensors offer a means to detect and quantify biophysical parameters, such as temperature, pH, radicals, magnetic fields, electric fields, and mechanical stress, with nanoscale spatial resolution and exceptional sensitivity^20^. Among these, fluorescent nanodiamonds (FNDs) hosting nitrogen-vacancy (NV) centers convert the state of an optically addressable electronic spin into stable fluorescence signals without photobleaching and blinking^20–23^. Using optically detected magnetic resonance (ODMR), thermometry is enabled by extracting the temperature-dependent axial zero-field splitting parameter *D* near the NV resonance frequency at ∼2,870 MHz, thereby providing a quantitative thermometer compatible with optical readout and free of cross-talk with other parameters^21,24–26^. In this study, we present a lensless fiber-optic quantum-sensing endomicroscopy platform that enables cellular-resolution thermometry in the mouse deep brain. This endomicroscopy can be used to dissect how deviations in biophysical parameters modulate deep-brain function.

## Results

### Setup of lensless quantum-sensing endomicroscopy

We first built a lensless fiber-optic quantum-sensing endomicroscopy system that combines wide-field fluorescence collection with local microwave excitation (Fig. 1). A copper microwire (50-μm diameter) was coupled along the distal edge of the 2.0-m imaging fiber bundle, in which 30,000 ± 3,000 individual quasi-single-mode glass fiber cores are assembled into a circular array (total diameter of 650 ± 30 μm, image-circle diameter of 600 ± 30 μm, and a 3.0 μm aperture at each fiber entrance). Therefore, the field of view (FOV) of imaging is 0.28 mm^2^. This microwire-coupled distal edge placed just above the FNDs delivered microwave (MW) irradiation at frequencies near the electron spin resonance of the NV centers. The proximal edge of the fiber bundle was coupled to a ×20/0.50 NA objective of an upright microscope. This system employed a lensless fiber bundle and does not require a GRIN lens^27–29^. Fluorescence imaging was performed using a 528 nm laser-diode light engine as the excitation source and a qCMOS camera as the detector. The laser was focused by the objective at the proximal edge of the fiber bundle. The sample temperature was maintained within ± 0.2 ℃ during each ODMR recording period by controlling a stage-top incubator in the *in vitro* experiments and the air conditioner in the *in vivo* experiments. Fluorescence intensities of the FNDs were recorded while sweeping the MW frequency from 2,850 to 2,890 MHz across 101 frequency points (Fig. 2a). We also adapted our previously established selective-imaging method which, owing to lock-in detection, is noise-tolerant and yields high signal-to-noise ratio (SNR) and signal-to-background ratio^30–32^. In this method, FND images are acquired with and without MW irradiation at each frequency, and the ratio of their fluorescence intensities is calculated for each frequency. Furthermore, one ODMR spectrum acquired with 101 frequency points was averaged over 30 accumulations. The ODMR spectra were fitted with a double-Lorentzian model^33^ using non-linear least squares to determine the zero-field splitting parameter *D*. We inserted the microwire-coupled imaging fiber-bundle and a thin thermocouple-type thermometer into an agarose gel mimicking mouse brain to a depth of 5 mm and confirmed that 9 dBm of MW irradiation did not mediate any detectable temperature increase during measurement.

**Fig. 1.**
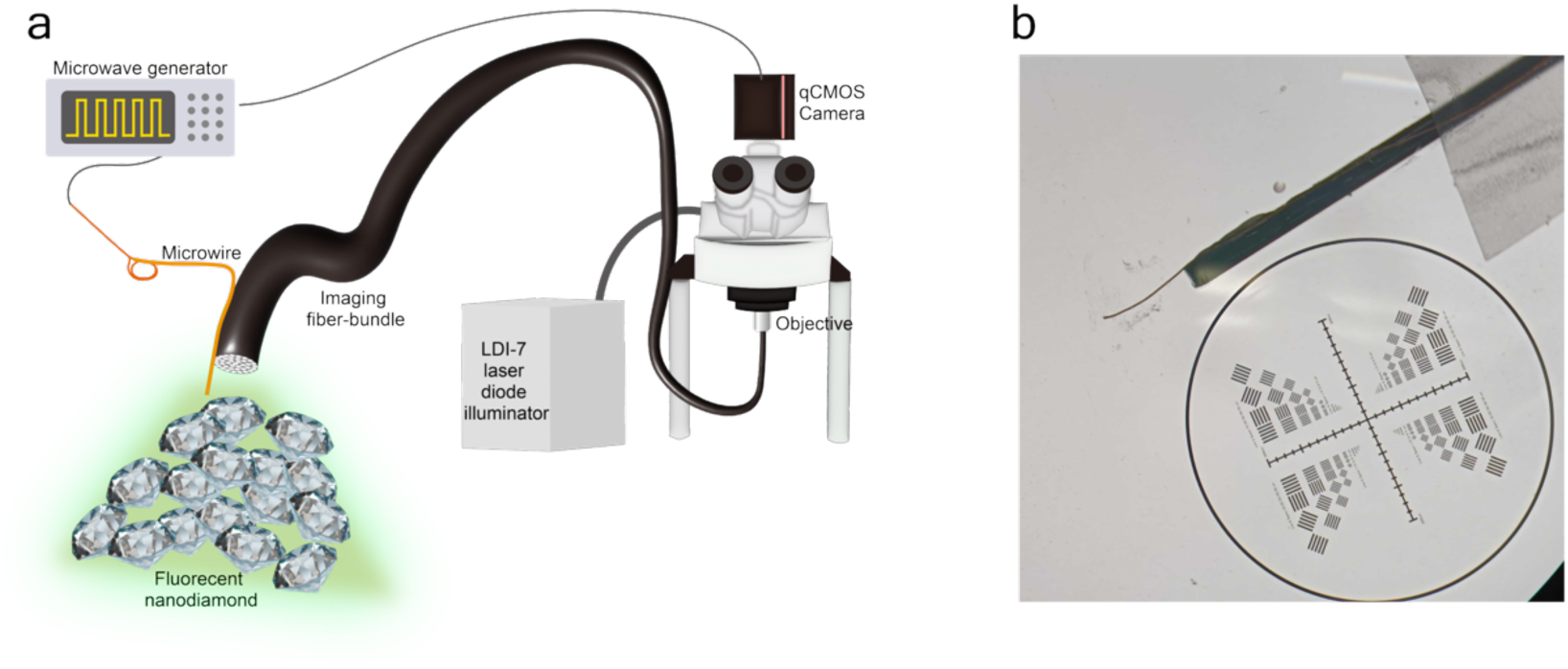
Setup of quantum-sensing endomicroscopy. **a**, Schematic diagram and photograph of the fiber-optic quantum-sensing endomicroscopy showing excitation/collection optics and microwire coupled to the imaging fiber. **b**, Close-up view of the distal tip of a microwire-coupled fiber bundle.

**Fig. 2.**
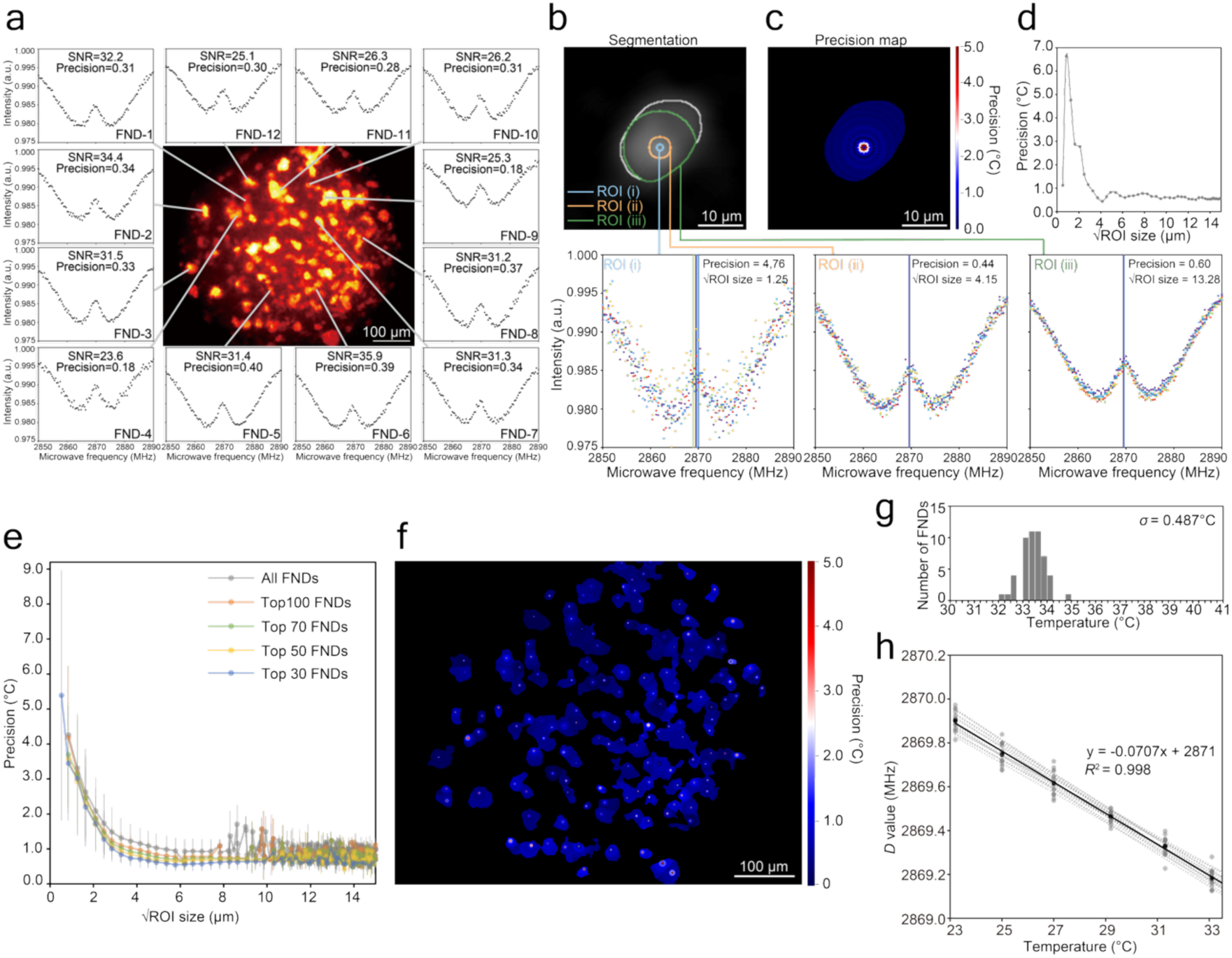
*In vitro* evaluation of a quantum-sensing endomicroscopy. **a**, Cladding-removed image of FNDs acquired via the fiber bundle, with representative ODMR spectra, SNRs, and precision values. **b–e**, Determination of the ROI size required to ensure measurement precision for all 146 FNDs. The initial full ROI (white line) was segmented and progressively reduced toward the centroid to generate a series of ROIs (green, orange, light blue). For each ROI, the ODMR spectra from six repeated measurements (shown as differently colored points) and the corresponding precision were computed. Fitted *D* values are indicated near 2,870 MHz to reflect precision. **c**, As the ROI was updated, precision was iteratively remapped and overlaid. Color bar indicates precision. Dependence of precision on square root of ROI size is shown for a representative FND (**d**) and the means across the top 30, 50, 70 and 100 FNDs ranked by precision among 146 FNDs (**e**). **f**, Precision map for all FNDs. Color bar represents temperature-measurement precision. **g**, Temperature variation histogram for the top 50 FNDs. **h**, Mean and individual accuracies for the top 15 FNDs ranked by the *R*^2^ of their calibration lines among 146 FNDs. Calibration lines for the mean across 15 FNDs (black) and individual FNDs (gray) are shown.

### Evaluation of a quantum-sensing endomicroscopy

We first evaluated, *in vitro*, the SNR of the ODMR spectra, the precision of the temperature measurements, and their accuracy^34^. SNR was defined as the ODMR contrast divided by the mean absolute fitting residual^35^ (Extended Data Fig. 1). Precision was quantified as the standard deviation of the fitted zero-field-splitting parameter *D* across six repeated measurements and subsequently converted to temperature using the calibration slope of *dD* / *dT* obtained from the accuracy analysis. In this framework, precision captures the repeatability of the measurement; a highly precise thermometer yields approximately identical readings under identical conditions. Accuracy was defined as the correspondence between temperature changes measured by the FNDs and those obtained using an on-sample thermocouple-type thermometer according to the calibration line.

Before imaging, a suspension of FNDs in PBS diluted two-fold with water was drop-cast onto a coverslip and air-dried. The distal end of the microwire-coupled fiber-bundle was then brought into contact with the dried film, enabling FNDs to attach to the bundle through passive adsorption and without any adhesive. Once attached, the FNDs remained stably bound to the fiber bundle even during insertion into the mouse brain. Raw images of *in vitro* FND samples acquired through the fiber bundle exhibited a characteristic honeycomb-like pattern arising from the negative space, largely occupied by cladding, between individual optical fibers over the fiber core (Extended Data Fig. 2a). We removed cladding contributions by applying a frequency filter^36^ (Extended Data Fig. 2b), identified FND regions of interest (ROIs), and extracted ODMR spectra by sweeping the microwave frequency. Representative ROIs, with or without cladding removal, produced comparable SNR values in the ODMR spectra (Extended Data Fig. 2c). Accordingly, all subsequent evaluations were performed using the cladding-removed images as shown in Fig. 2a.

In general, smaller FNDs or ROIs yield lower fluorescence intensities and greater variance in the fitted *D* values, thereby reducing temperature precision, whereas larger FNDs or ROIs improve precision at the expense of spatial resolution. To determine whether our fiber-optic endomicroscopy achieves cellular-level resolution with adequate precision, we systematically quantified the relationship between ROI size and measurement precision (Fig. 2b–e). Each FND was first segmented to generate an initial full ROI, and the ROI was then progressively reduced toward the centroid to produce a series of nested ROIs (Fig. 2b). For each ROI, we computed the ODMR spectrum and its associated precision (Fig. 2b, ROI(i)–(iii); Fig. 2c,d). ROI size and corresponding precision were obtained for all FNDs (*n* = 146). Across the top 50 FNDs ranked by precision, the mean ROI size and mean precision were 4.88 μm and 0.49 ℃, respectively (Fig. 2e). The minimum (best) precision observed among the 146 FNDs was 0.18 ℃. The precision map in Fig. 2f demonstrates that our fiber-optic endomicroscopy can resolve temperature across multiple local regions at cellular resolution with sufficient precision. This high level of precision was further reflected in the minimal temperature variation across the top 50 FNDs (Fig. 2g). We also verified that precision estimates for 20 FNDs obtained through Monte Carlo simulation^31,34^ closely matched those from repeated *in vitro* measurements, with a mean difference of 0.0075 ± 0.104 ℃ (mean ± s.d.; Extended Data Fig. 3). This agreement supports the use of Monte Carlo-derived precision estimates for subsequent *in vivo* experiments, where repeated measurements may be influenced by physiological temperature fluctuations. To further assess the accuracy of our fiber-optic endomicroscopy, we generated a calibration line with low residuals (−5.7 × 10⁻¹³ ± 0.045 MHz; mean ± s.d.), confirming that the fiber-optic thermometry accurately reports temperature (Fig. 2h). Together, these results demonstrate that our endomicroscopy captures temperature and its deviations with enough precision and accuracy *in vitro*.

### *In vivo* proof-of-principle experiment conducted in the deep brain regions of mice

We next performed an *in vivo* proof-of-principle experiment in head-fixed mice (Fig. 3a,b). To determine the absolute temperature, we first measured the mean calibration line of 15 FNDs adhered to the distal end of the fiber bundle in a temperature-controlled chamber *in vitro* before the *in vivo* experiment (Extended Data Fig. 4). We also estimated temperature precision for all FNDs using a Monte Carlo simulation. The same FND-coated, microwire-coupled fiber bundle was then inserted through the cortical surface at bregma. Using bregma as the stereotaxic reference, we positioned the distal end of the fiber bundle and lowered the probe in 1-mm increments, acquiring measurements down to the ventromedial hypothalamic nucleus (VMH) at a depth of 5 mm (Fig. 3c,d). Because the same FNDs were carried along the tract, primarily the same FNDs were measured at each depth. Our quantum-sensing endomicroscopy enabled acquisition of ODMR spectra at all depths examined (Fig. 3e). Across the entire depth range, the SNRs and precision values of the FNDs were comparable to those obtained *in vitro* (Fig. 2a), with SNRs of 29.5 ± 3.86 (mean ± s.d.) *in vitro* and 32.5 ± 2.14 *in vivo*, and precisions of 0.31 ± 0.07 ℃ *in vitro* and 0.72 ± 0.04 ℃ *in vivo*. We then performed temperature mapping at each depth using 72 FNDs with Monte Carlo-estimated precision < 1 ℃ at the 5-mm depth (Fig. 3f). From these maps, we found that the mean FND-reported temperatures were influenced by ambient temperature (24–26 ℃) from 0–2 mm depth and approached core body temperature below ∼3 mm (Fig. 3g). Furthermore, comparison of the temperature-variation histograms from the *in vitro* (Fig. 2g) and *in vivo* experiments (Fig. 3h) indicates that even at 5 mm depth, the temperature distribution remained heterogeneous rather than uniform, with variations of 0.49 ℃ and 1.28 ℃ for the *in vitro* and *in vivo* experiments, respectively. We next mapped temperatures under awake and anesthetized conditions (Fig. 3i). Anesthesia induced reductions in deep-brain temperature of –4.18 ± 1.94 ℃ (mean ± s.d. across 66 FNDs), along with reductions in rectal temperature measured using a thermocouple thermometer, consistent with reduced metabolic heat production under anesthesia^37^ (Fig. 3j). In addition, the temperature-deviation map and histogram indicated that this reduction occurred in a spatially nonuniform manner (Fig. 3k,l).

**Fig. 3.**
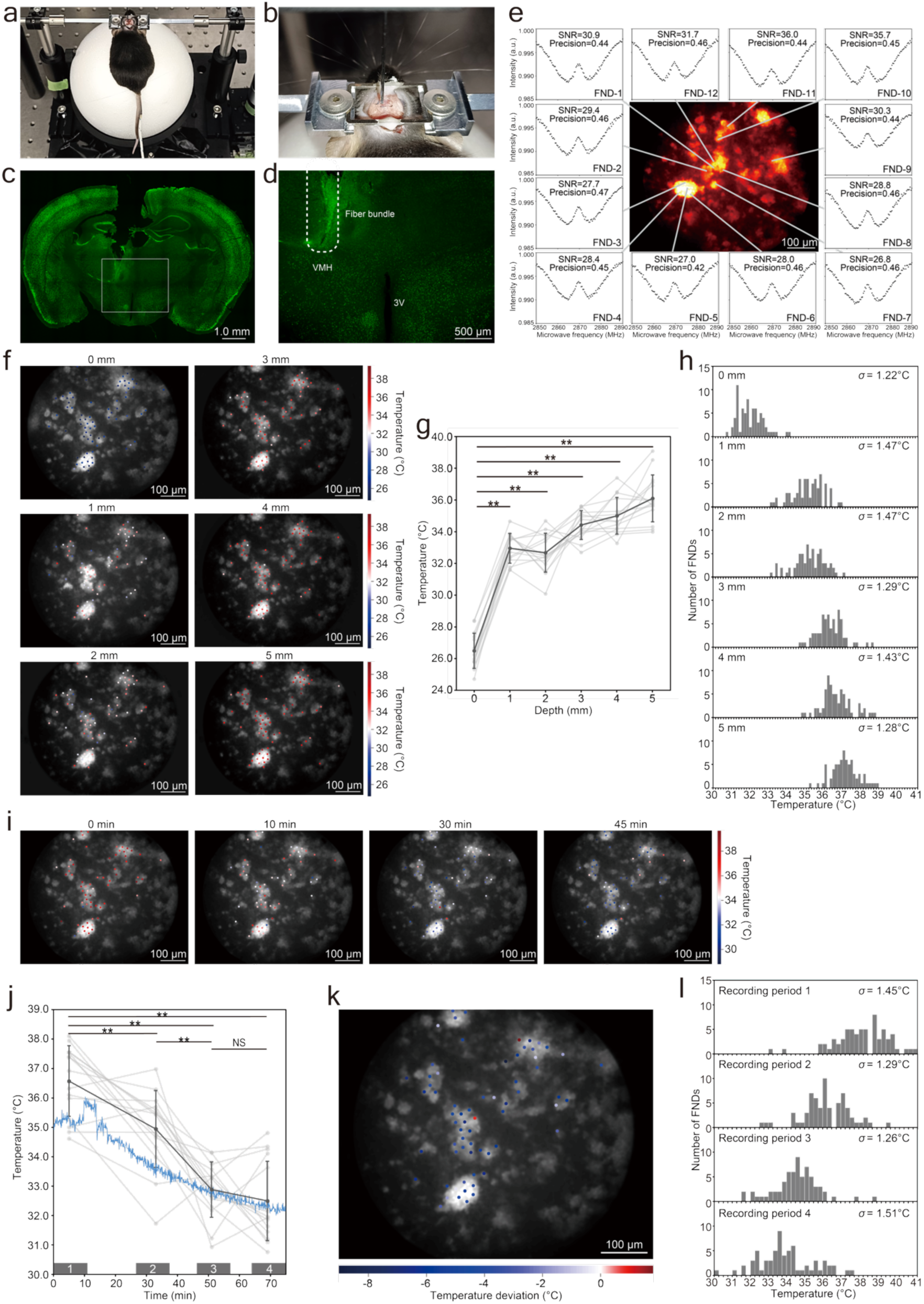
Deep-brain, cellular-resolution thermometry in head-fixed mouse. **a,b**, Photographs of a head-fixed mouse on a custom-made treadmill and a close-up of fiber-bundle-implanted head. **c,d** Brain section from the region imaged by quantum-sensing endomicroscopy at a depth of 5 mm. Neurons were labeled via NeuN immunostaining after imaging. Regions corresponding to ventromedial hypothalamic nucleus (VMH) and third ventricle (3V) in (**c**) are shown at higher magnification in (**d**). **e**, ODMR spectra of the top 12 FNDs ranked by precision among 185 FNDs at a depth of 5 mm, with corresponding SNRs and precisions. **f**, *In vivo* temperature mapping at the indicated depths. Measurements were performed at room temperature (24–26 ℃). At each depth, 72 FNDs with Monte Carlo-estimated precision < 1 ℃ and ROI size < 10 μm (from data at 5 mm depth) are mapped. Color bar indicates absolute temperature for all FNDs, estimated using individual calibration slopes (Extended Data Fig. 4). **g**, Temperature at each brain depth. Mean temperature of the top 15 FNDs ranked by Monte Carlo-estimated precision among 185 FNDs (black) and their individual temperatures (gray) are shown. Statistical analysis was performed using Student’s *t*-test. \*\**P* < 0.01. Error bars, ± s.d. **h**, Temperature-variation histogram based on temperatures of the 72 FNDs. **i**, Temperature maps acquired before and during isoflurane anesthesia. A total of 66 FNDs with Monte Carlo-estimated precision < 1 ℃ and ROI size < 10 μm are mapped for each period. Color bar indicates absolute temperature estimated from calibration slopes. **j**, Temperature traces during anesthesia measured with the fiber bundle in VMH at 5 mm brain depth and with a thermocouple-type thermometer in the rectum. Mean temperature trace of the top 15 FNDs (black), the rectal temperature trace (light blue), and individual FND traces (gray) are shown. Top 15 FNDs were selected by Monte Carlo-estimated precision among 137 FNDs. Bars mark one pre-anesthesia period (0–10 min) and three anesthesia periods (28–38 min, 46–56 min, 64–74 min). Statistical analysis was performed using Student’s *t*-test. \*\**P* < 0.01; NS, not significant. Error bars, ± s.d. **k**, Temperature-deviation map comparing recording periods during awake (0–10 min) and anesthetized (64–74 min) states in VMH. Color bar indicates temperature difference. **l**, Temperature-variation histogram created from temperatures of the 66 FNDs.

We finally examined thermometry in freely behaving mice (Fig. 4). The mouse was allowed to move freely in its home cage without disrupting the FOV (Fig. 4a–c). Temperature traces based on the mean of 15 FNDs across four 10-min recording periods indicated that, in VMH, temperature appeared macroscopically stable under steady-state conditions (Fig. 4d). However, temperature map and temperature-deviation map comparing the first and fourth recording periods showed that both the spatial distribution and the temporal change were heterogeneous rather than uniform, with a maximum spatial temperature difference of 7.39 ℃ in distribution and a maximum temporal change of 4.12 ℃ over 74 min (Fig. 4e,f). The temperature-variation histogram likewise supported this *in vivo*-specific heterogeneity (Fig. 4g). Next, we applied a tail pinch as a nociceptive stimulus^38^, which elicited vigorous locomotion (Fig. 4h,i). Despite this movement, we successfully acquired ODMR spectra and generated temperature maps and temperature-variation histograms (Fig. 4j,k). Together, these proof-of-principle experiments demonstrate that our endomicroscopy enables deep-brain, cellular-resolution thermometry even during vigorous movement.

**Fig. 4.**
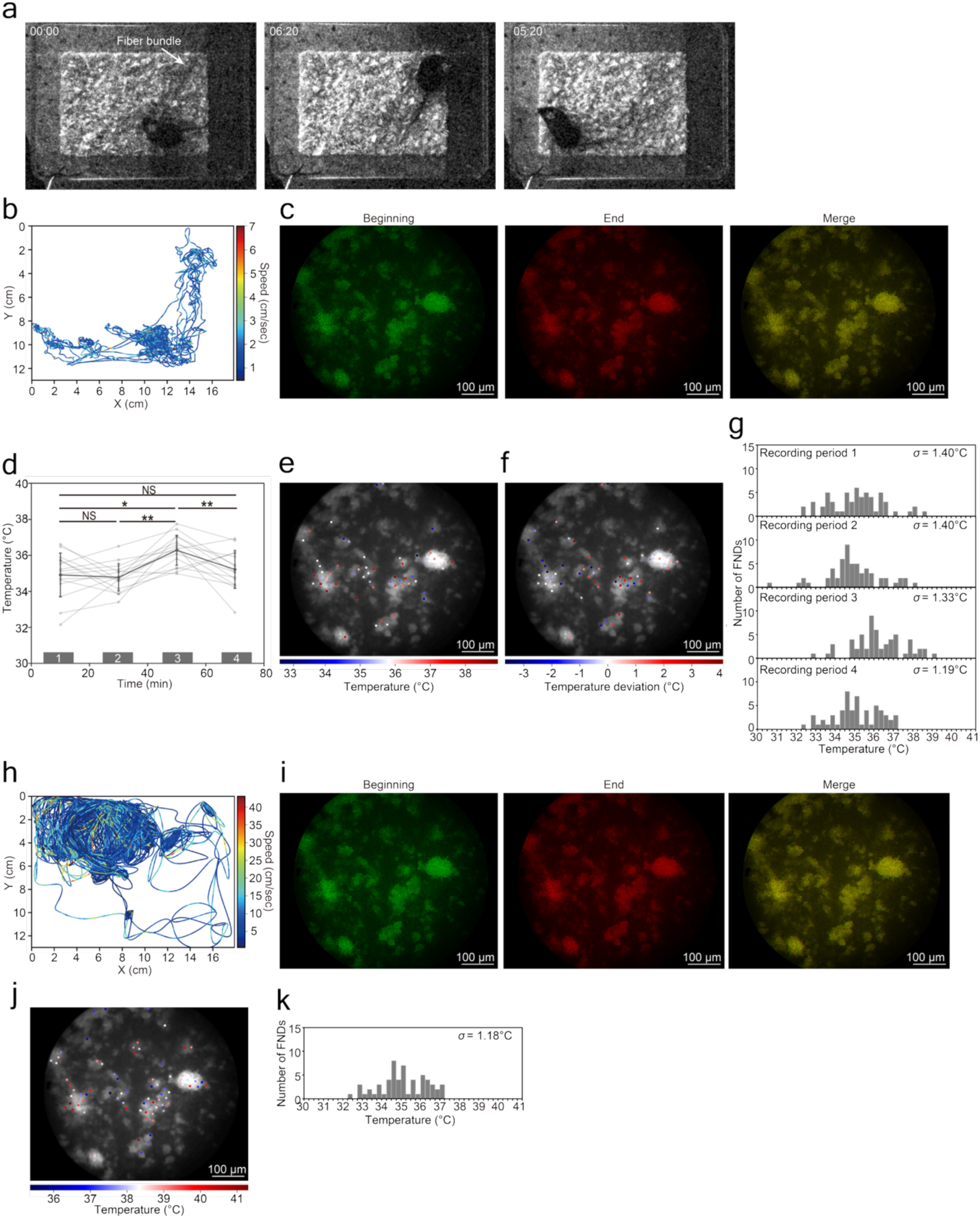
Deep-brain cellular-resolution thermometry in a freely behaving mouse. **a,b**, Representative images of a freely moving mouse in its home cage with a fiber bundle implanted at a brain depth of 5 mm, together with the corresponding trajectory over a 10-min recording period. **c**, FOVs acquired at the beginning and end of the recording period and their overlay, illustrating FOV stability. Total displacement over the entire period was < 1.04 μm. **d**, Temperature traces measured at 5 mm depth in a freely moving mouse across four recording periods. Mean temperature of the top 15 FNDs ranked by precision among 137 FNDs (black) and individual FND temperatures (gray) are shown. Statistical analysis was performed using Student’s *t*-test. \*\**P* < 0.01; \**P* < 0.05; NS, not significant. Error bars, ± s.d. **e**, Temperature map at 5 mm depth from the third recording period. Color bar represents temperature. A total of 58 FNDs with Monte Carlo-estimated precision < 1 ℃ and ROI size < 10 μm are mapped. **f**, Temperature-deviation map at 5 mm depth, comparing the first and fourth recording periods. **g**, Temperature-variation histogram created from temperatures of 58 FNDs. Color bar represents temperature deviation. **h**, Trajectory of the mouse in response to a nociceptive stimulus during the recording period. **i**, FOVs at the beginning and end of the period and their overlay. Total displacement over the entire period is < 1.61 μm. **j**, Temperature map recorded in VMH. A total of 53 FNDs are mapped with Monte Carlo-estimated precision < 1 ℃ and ROI size < 15 μm. **k**, Temperature-variation histogram created from temperatures of 53 FNDs.

## Discussion

We introduced a fiber-optic quantum-sensing endomicroscopy that maps temperature in the mouse deep brain by reading ODMR spectra from multiple FNDs through a microwire-coupled imaging fiber bundle. This approach leveraged the temperature-dependent shift of the NV-center zero-field splitting to extract temperature deviations and achieved mapping across several deep-brain regions down to 5 mm. A practical advantage of our endomicroscopy is the exceptional photostability of FND, which enabled repeated or prolonged acquisitions without signal loss^39^. A second advantage is that FNDs are exogenous probes and therefore require no protein expression or maturation period, unlike fluorescent-protein reporters that often need a few weeks for *in vivo* expression^40^. Finally, because our system relies on a lensless fiber bundle rather than a GRIN lens, it avoids the intrinsic aberrations^19^.

Our current system is limited in temporal resolution because many frequency sweeps and substantial frame averaging are needed to obtain robust ODMR fits. This limitation can be mitigated by sampling only a few informative frequencies around the resonance, using adaptive acquisition that terminates averaging once a target precision is reached, and modestly increasing photon rates through improved collection optics or brighter nanodiamonds to shorten camera exposure. A second limitation is that the present system produces only a two-dimensional image, since a single image plane is formed at the bundle face and depth information is integrated along the line of sight. To extend this to three-dimensional imaging without time-consuming mechanical axial scans, we can adopt a computational light-field strategy in which the mode structure within each fiber core is used to recover the spatio-angular information of the captured rays as the light field, thereby enabling three-dimensional reconstruction^41^. A third limitation is that, at present, most FNDs likely reside extracellularly rather than inside identified cells. This issue can be addressed by targeted intracellular delivery using tailored surface chemistry^39,42–44^ and peptide or antibody conjugation for receptor-mediated uptake^45^ .

By combining cellular-scale temperature maps with modern neural circuit readouts and atlases, this technology can test how spatially confined thermal microgradients regulate hypothalamic and circadian neural circuitry *in vivo* and during behavior. The deeper access compared with two-photon methods suggests that synergy with fiber imaging of GCaMP or three-photon optical recordings can provide complementary population-level signals of neural activities^16,46^. More broadly, NV centers enable multiparameter quantum sensing of magnetic and electric fields as well as temperature and pH, opening opportunities for co-localized readouts of metabolic heat, neuromodulator dynamics, and endogenous fields in health and disease^23,47^. Prior *in vivo* quantum thermometry in small organisms such as *C. elegans*^25^ has established feasibility for real-time biology, and adapting the present platform to miniaturized probes and faster spectroscopic protocols could extend such studies to freely behaving mammals. Finally, because optical fiber bundles are widely used in the medical field, our approach could be adapted to diagnostic applications, such as in fiber-bundle biopsy.

## Methods

### Optical setup

The proximal end of 2 m multicore optical fiber bundle (Fujikura, FIGH-30-650S) was coupled to a ×20/0.50 NA objective (Olympus, UPlanFI) on a BX51 upright microscope (Olympus). The distal end of the fiber bundle was coupled to a microwire (thin copper wire; 50 μm). Laser light from the LDI-7 (89North) was delivered to the focal plane of the microscope objective, where the proximal fiber end was positioned, transmitted through the bundle, and delivered onto the FNDs. Returning fluorescence was imaged onto the distal fiber end, transmitted through the bundle and microscope, and reimaged on a qCMOS camera (Hamamatsu Photonics, ORCA-quest). The fiber bundle comprised 30,000 individual fiber cores distributed across a 600-μm diameter imaging area. The BX51 upright microscope was equipped with a 565 nm dichroic mirror (Olympus, U-MWIG2) and a 650 nm long-pass filter (Thorlabs, FELH0650). Data were acquired using HCImage software (Hamamatsu Photonics) on an Intel Xeon workstation (W-2133, 3.60 GHz; 512 GB RAM).

### Preparation of FND

Commercial nanodiamonds (Micron+ MDA 0–0.1 μm, Element Six) were irradiated with electrons at 2 MeV to a fluence of 4 × 10¹⁸ cm⁻². The samples were annealed at 800 °C for 2 h under reduced pressure to form NV centers, followed by air oxidation at 550 °C for 2 h to remove surface graphitic carbon. To remove residual surface contaminants and enrich surface carboxyl groups, the particles were treated in H₂SO₄:HNO₃ (9:1, v/v) at 70 °C for 3 days. The suspensions were then centrifuged (12,000 × g, 10 min), the supernatants were discarded, and the pellets were washed thrice with Milli-Q water. Next, the pellets were stirred in 0.1 M NaOH at 90 °C for 2 h and centrifuged, washed with Milli-Q water, and then stirred in 0.1 M HCl at 90 °C for 2 h. Finally, the particles were washed thrice with Milli-Q water, yielding FNDs.

### ODMR measurement

Before all imaging, including *in vivo* experiments, a suspension of FNDs in PBS diluted two-fold with water was drop-cast onto a coverslip and air-dried. The distal end of the microwire-coupled fiber bundle was then brought into contact with the dried film to attach FNDs to the bundle by passive adsorption. The MW pulses were generated by an MW generator (Keysight, N5181A MXG) and amplified using linear MW power amplifiers (Mini Circuits, ZHL-16W-43-S+). Synchronization between MW irradiation and image acquisition was achieved using a function generator (NF Corporation, WF1974). ODMR spectra were recorded across a range of MW frequencies by calculating the ratio of fluorescence intensities with and without MW irradiation. The sample temperature was controlled using a stage-top incubator (Tokai Hit, STXGC-KRiK-SET; precision, ± 0.3 ℃). During accuracy measurements, the sample temperature was monitored using a thermocouple-type thermometer (Graph Tech, GL10-TK and Twtade, 768894228954; resolution, 0.06 ℃). The MW frequency was digitally swept from 2,850 to 2,890 MHz in 101 frequency points. The qCMOS camera exposure time was set to 100 ms. For each MW frequency, a set of images was accumulated 30 times across all experiments.

### ODMR spectrum and *D* value extraction

The captured image stack, consisting of fluorescence images at each MW frequency, was processed using a custom Python script. The ROI for each FND was determined as described in the analysis of the dependence of precision on ROI size. The fluorescence intensity as a function of MW frequency was extracted for each FND to generate ODMR spectra. To determine the zero-field splitting parameter *D*, the ODMR spectra were fitted using a double-Lorentzian model with nonlinear least squares. In the presence of no external magnetic field, the two dips *f*± were estimated from the fits, and the *D*-value was calculated as (*f*++ *f*−) / 2.

### Monte Carlo simulation to estimate temperature measurement precision

We estimated the achievable temperature precision using a Monte Carlo simulation that models spectral variability by incorporating measured noise characteristics^48^. First, the experimentally obtained ODMR spectrum was used as the reference after averaging the signal across 30 accumulations. To better capture the variability of the original spectrum, the reference spectrum was fitted with a Lorentzian–Gaussian–Lorentzian model, which provided a superior fit compared with a double-Lorentzian model. We then calculated the noise level and its standard deviation from the reference spectrum and used these values as inputs to the simulation. Synthetic spectra sampled at 101 frequency points between 2,850 and 2,890 MHz were generated by adding Gaussian noise to the fluorescence intensity to mimic experimental fluctuations. Each simulated spectrum was fitted with a double-Lorentzian model to extract the zero-field splitting parameter *D*. The standard deviation of *D* values across 300 simulated spectra was then calculated. For each FND, this standard deviation was converted to temperature precision using the calibration slope determined *in vitro* (Extended Data Fig. 4).

### Analysis of the dependence of precision on ROI size

All analyses were performed in Python. The FND image was binarized with OpenCV, and individual FNDs were segmented to obtain an initial full ROI using cv2.findContours. The centroid of each FND was identified using SciPy’s maximum_filter. The ROI for each FND was then progressively reduced toward the centroid, and for each ROI, we computed the ODMR spectrum and its precision. Precisions for *in vitro* and *in vivo* FNDs were obtained from six repeated measurement datasets and Monte Carlo simulations, using previously reported *dD* / *dT*, values of –74 kHz / ℃^26,49^, respectively. Plotting precision against the square root of ROI size revealed ROI size that yielded the minimum (best) precision. This ROI size and its corresponding precision were assigned to each FND. Finally, the precision assigned to each FND was converted using the mean slope of the calibration lines for the top 15 FNDs ranked by experimentally determined *R*^2^ values (as shown in Fig. 2h and Extended Data Fig. 4; *dD* / *dT* = –70.7 and –57.1 kHz / ℃ for the *in vitro* and *in vivo* experiments, respectively).

### *In vivo* quantum-sensing endomicroscopic imaging in mice

All procedures conformed to the guidelines or animal care established by Hiroshima University and were approved by the Hiroshima University Animal Experiment Committees. Male C57BL/6 mice (aged seven weeks, CLEA Japan, Inc) were used without randomization or blinding. All mice were housed with ad libitum access to food and water in an air-conditioned room at 23 ℃ with a 12-h light/12-h dark cycle (lights at 7 a.m.). The relative humidity was 50%. The animals were anesthetized with a mixture of air and 2% isoflurane.

The FND-attached fiber bundle was inserted toward the hypothalamus from a stereotaxic entry point (from bregma: anteroposterior, −1.5 mm; mediolateral, +0.3 mm; and from dura: dorsoventral, −5.0 mm) in a head-fixed mouse. The head-fixed mouse was placed on a custom-made treadmill (Fig. 3a,b) using (Narishige, MAG-A) and (Narishige, CF-10), after which imaging was conducted. For freely behaving experiments, the mouse with the preinserted fiber bundle was transferred to a home cage (CLEA Japan, 172 mm (W) × 240 mm (D) × 129 mm (H), CL-0103-1), and imaging was carried out during free movement. Tail pinching was applied using a clamp (As One, No. 23, 48 mm)^38^ .

### Immunohistochemistry

The following procedures were performed at room temperature unless stated otherwise. Mouse was deeply anesthetized and transcardially perfused with 4% formaldehyde in 0.1 M PB, pH 7.4. The brain was removed, postfixed in the fixative at 4°C for 2-3 h, and then cryoprotected in a 30% sucrose solution overnight. The tissue was cut into 30 mm-thick frontal sections on a freezing microtome. For detection of neuronal cell bodies, section was incubated overnight with an anti-NeuN mouse antibody (1:1000, MAB377, Merck Millipore). After a rinse, the sections were incubated for 1 h with an Alexa488-conjugated goat antibody to mouse IgG (5 µg/ml; A11034, Fisher Scientific). Stained sections were mounted on glass slides and covered with a mounting medium (50% glycerol/50% PBS containing 2.5% triethylenediamine). The sections were observed under a fluorescence microscope (Eclipse 80i, Nikon), or a confocal laser scanning microscope (TCS SP8, Leica).

## Acknowledgements

We thank Marie Hirose, Nozomi Tominaga, and Aoi Yoshimatsu for technical assistance and Prof. Kazuhiro Nakamura (Nagoya University Graduate School of Medicine) for kindly sharing his laboratory’s resources. This work was supported by JST-FOREST Program (JPMJFR214R to T.S. and JPMJFR2428 to S.S.), JSPS KAKENHI Scientific Research (A) (24H00577 to T.S.), Grant-in-Aid for Challenging Research (Pioneering) (23K17405 to T.S.), Grant-in-Aid for Transformative Research Areas B (JP23H03845 to T.S. and JP23H03844 to N.K.), Grant-in-Aid for JSPS Fellows (24KJ1745 to R.I.), JPSP program for Forming Japan’s Peak Research Universities (J-PEAKS, JPJS00420230011).

## Author contributions

T.S. conceived the project with R.I. R.I., Y.N. performed experiments with help from N.K. S.S., H.A., and T.A. prepared FND. T.S. supervised the project and wrote the manuscript with R.I. and Y.N. All authors have approved the final version of the manuscript.

## Data availability

All data supporting the findings in this study are available from the corresponding authors on reasonable request.

## Supplementary Information

**Extended Data Fig. 1.**
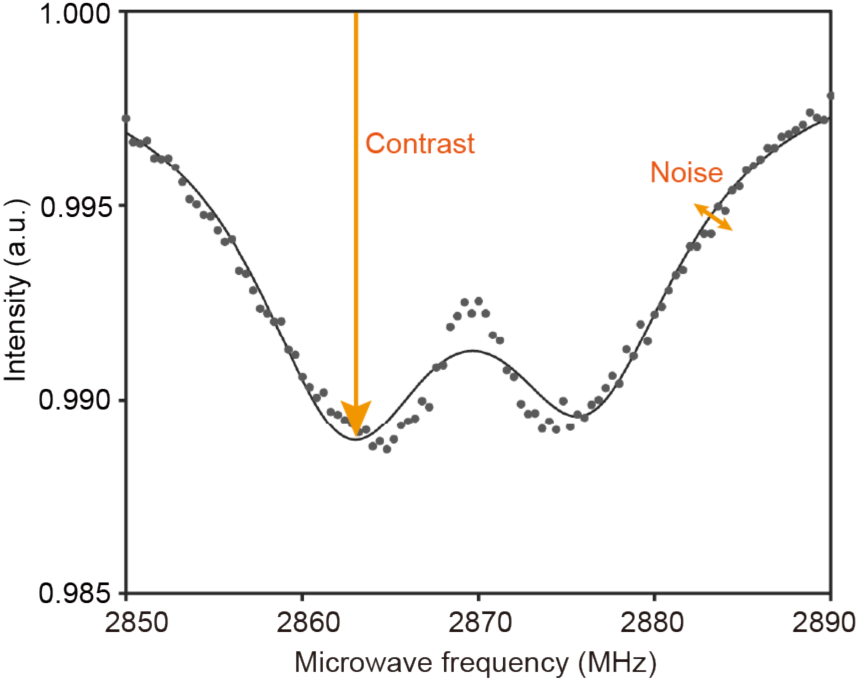
SNR definition for ODMR spectrum. SNR was characterized as the ODMR contrast divided by the mean absolute fitting residual.

**Extended Data Fig. 2.**
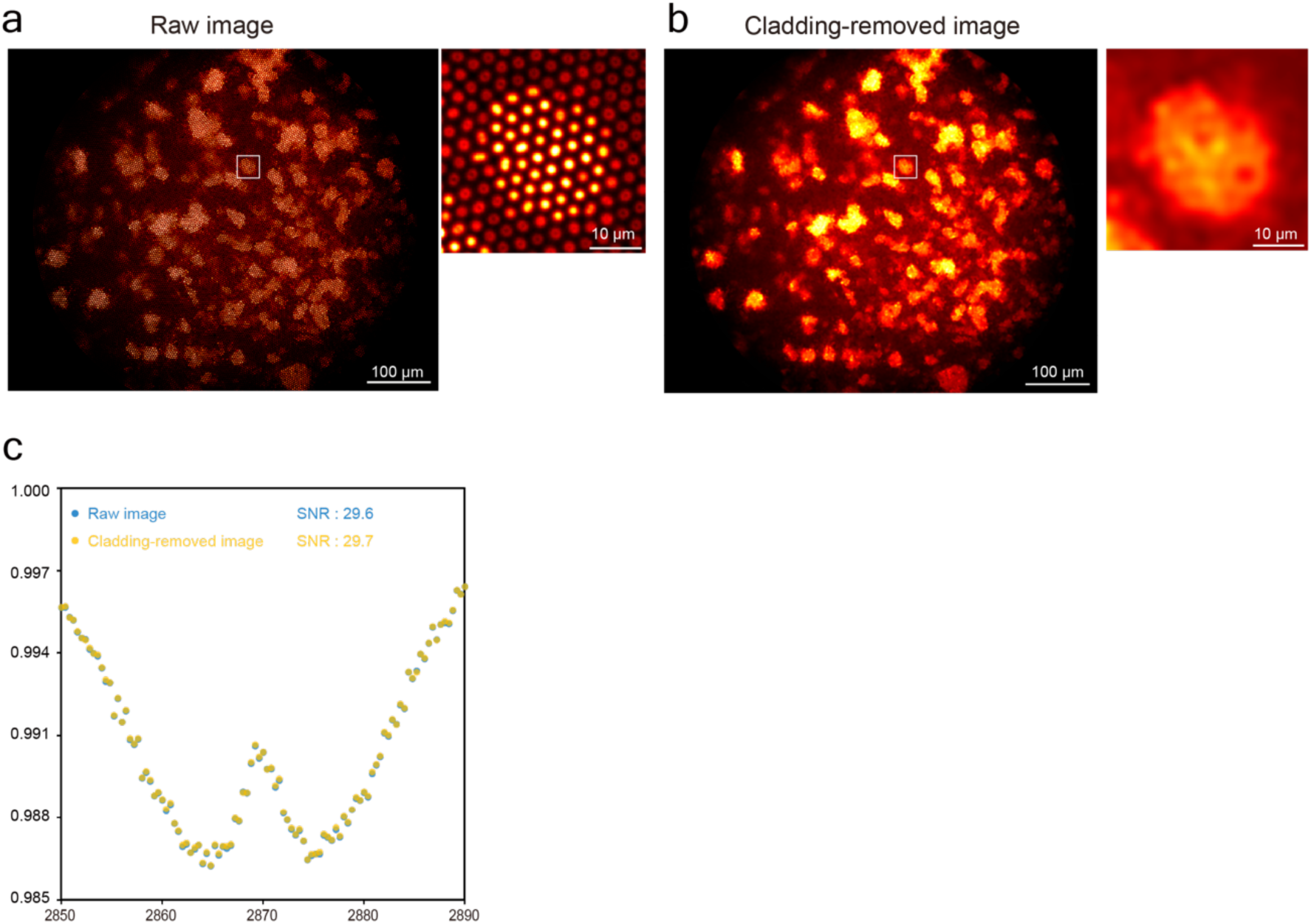
Effect of cladding removal on ODMR spectrum. **a,b**, Raw image (**a**) and cladding-removed image (**b**) of *in vitro* FND samples with extracted ODMR spectra and SNRs (**c**).

**Extended Data Fig. 3.**
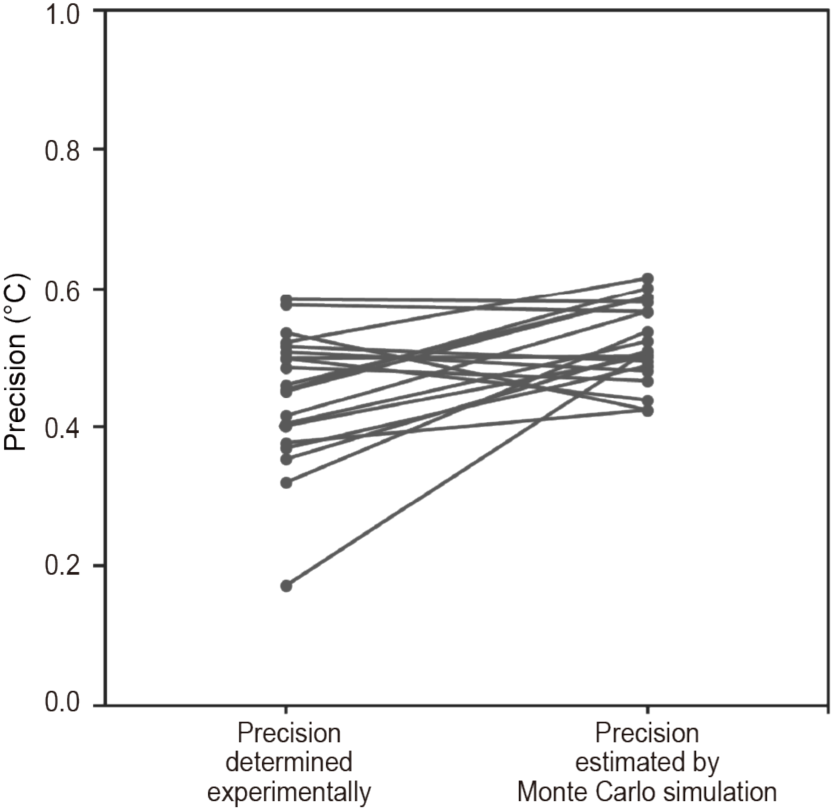
Precision estimation of *in vitro* FNDs by Monte Carlo simulation. Precision values for 20 FNDs were determined based on six repeated measurements under identical *in vitro* temperature conditions or were estimated from a single ODMR spectrum via Monte Carlo simulation. Simulated precision closely matched that obtained from repeated *in vitro* measurements.

**Extended Data Fig. 4.**
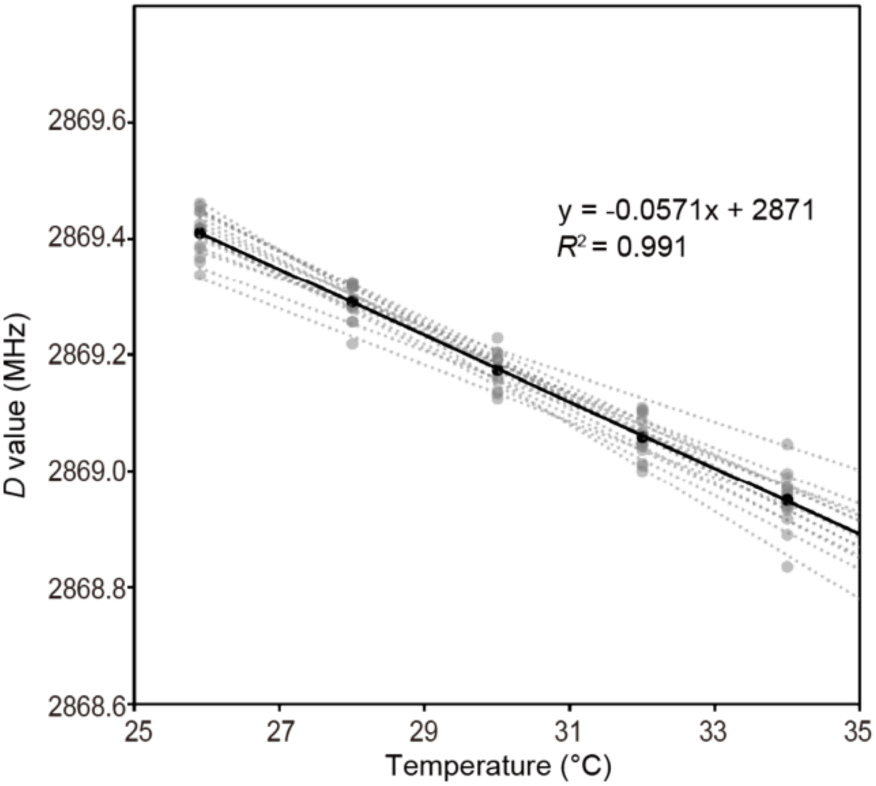
*In vitro* accuracy measurements of FNDs attached to the fiber bundle for implantation in mice. Mean and individual accuracies for 15 FNDs. Calibration lines for mean of 15 FNDs and individual FNDs are shown in black and in gray, respectively.

## References

1. Lin, Z., Xuan, Y., Zhang, Y., Zhou, Q. & Qiu, W. Hypothalamus and brainstem circuits in the regulation of glucose homeostasis. Am. J. Physiol.-Endocrinol. Metab. 328, E588–E598 (2025).

2. Morf, J. & Schibler, U. Body temperature cycles: Gatekeepers of circadian clocks. Cell Cycle 12, 539–540 (2013).

3. Tran, L. T. et al. Hypothalamic control of energy expenditure and thermogenesis. Exp. Mol. Med. 54, 358–369 (2022).

4. Hook, M. J. V. Temperature effects on synaptic transmission and neuronal function in the visual thalamus. PLoS ONE 15, e0232451 (2020).

5. Feng, Z., Saha, L., Dritsa, C., Wan, Q. & Glebov, O. O. Temperature-dependent structural plasticity of hippocampal synapses. Front. Cell. Neurosci. 16, 1009970 (2022).

6. Wang, H. et al. Brain temperature and its fundamental properties: a review for clinical neuroscientists. Front. Neurosci. 8, 307 (2014).

7. Xie, Y. et al. Temperature effects on the neuronal dynamics and Hamilton energy. *Chaos*, Solitons Fractals 195, 116325 (2025).

8. Conti, B. et al. Transgenic mice with a reduced core body temperature have an increased life span. 314, 825–828 (2006).

9. Kamm, G. B. et al. A synaptic temperature sensor for body cooling. Neuron 109, 3283–3297.e11 (2021).

10. Ziemann, A. E. et al. The Amygdala Is a Chemosensor that Detects Carbon Dioxide and Acidosis to Elicit Fear Behavior. Cell 139, 1012–1021 (2009).

11. Wemmie, J. A., Taugher, R. J. & Kreple, C. J. Acid-sensing ion channels in pain and disease. Nat. Rev. Neurosci. 14, 461–471 (2013).

12. Liu, Q. et al. Infrared thermography in clinical practice: a literature review. Eur. J. Méd. Res. 30, 33 (2025).

13. Blakley, S. et al. Fiber-Optic Quantum Thermometry with Germanium-Vacancy Centers in Diamond. ACS Photonics 6, 1690–1693 (2019).

14. Fedotov, I. V. et al. Fiber-based thermometry using optically detected magnetic resonance. Appl. Phys. Lett. 105, 261109 (2014).

15. Sui, K. et al. Temperature sensing of the brain enabled by directly inscribed Bragg gratings in CYTOP polymer optical fiber implants. Opt. Fiber Technol. 80, 103478 (2023).

16. Simpson, E. H. et al. Lights, fiber, action! A primer on in vivo fiber photometry. Neuron 112, 718–739 (2024).

17. Mattay, R. R. et al. MR Thermometry during Transcranial MR Imaging–Guided Focused Ultrasound Procedures: A Review. *Am*. J. Neuroradiol. 45, 1–8 (2023).

18. Takasaki, K., Abbasi-Asl, R. & Waters, J. Superficial Bound of the Depth Limit of Two-Photon Imaging in Mouse Brain. eNeuro 7, ENEURO.0255-19.2019 (2020).

19. Lee, W. M. & Yun, S. H. Adaptive aberration correction of GRIN lenses for confocal endomicroscopy. Opt. Lett. 36, 4608 (2011).

20. Aslam, N. et al. Quantum sensors for biomedical applications. Nat. Rev. Phys. 5, 157–169 (2023).

21. Kucsko, G. et al. Nanometre-scale thermometry in a living cell. Nature 500, 54–58 (2013).

22. Sotoma, S., Okita, H., Chuma, S. & Harada, Y. Quantum nanodiamonds for sensing of biological quantities: angle, temperature, and thermal conductivity. Biophysics Physicobiology e190034 (2022) doi:10.2142/biophysico.bppb-v19.0034.

23. Schirhagl, R., Chang, K., Loretz, M. & Degen, C. L. Nitrogen-Vacancy Centers in Diamond: Nanoscale Sensors for Physics and Biology. Phys. Chem. 65, 83–105 (2014).

24. Neumann, P. et al. High-Precision Nanoscale Temperature Sensing Using Single Defects in Diamond. Nano Letters (2013) doi:10.1021/nl401216y.

25. Fujiwara, M. et al. Real-time nanodiamond thermometry probing in vivo thermogenic responses. Sci Adv 6, eaba9636 (2020).

26. Acosta, V. M. et al. Temperature Dependence of the Nitrogen-Vacancy Magnetic Resonance in Diamond. Phys Rev Lett 104, 070801 (2010).

27. Badt, N. & Katz, O. Real-time holographic lensless micro-endoscopy through flexible fibers via fiber bundle distal holography. Nat. Commun. 13, 6055 (2022).

28. Song, P. et al. Ptycho-endoscopy on a lensless ultrathin fiber bundle tip. Light: Sci. Appl. 13, 168 (2024).

29. Sun, J. et al. Quantitative phase imaging through an ultra-thin lensless fiber endoscope. Light: Sci. Appl. 11, 204 (2022).

30. Igarashi, R. et al. Real-Time Background-Free Selective Imaging of Fluorescent Nanodiamonds in Vivo. Nano Lett 12, 5726–5732 (2012).

31. Yanagi, T. et al. All-Optical Wide-Field Selective Imaging of Fluorescent Nanodiamonds in Cells, In Vivo and Ex Vivo. Acs Nano 15, 12869–12879 (2021).

32. Miller, B. S. et al. Spin-enhanced nanodiamond biosensing for ultrasensitive diagnostics. Nature 587, 588–593 (2020).

33. Shiraya, K. et al. Hybrid nanosensors of carbon quantum dots and fluorescent nanodiamonds: Ratiometric thermometry and multicolor sensing. Carbon 242, 120457 (2025).

34. Yanagi, T., Kaminaga, K., Kada, W., Hanaizumi, O. & Igarashi, R. Optimization of Wide-Field ODMR Measurements Using Fluorescent Nanodiamonds to Improve Temperature Determination Accuracy. Nanomaterials 10, 2282 (2020).

35. Fujisaku, T., So, F. T. K., Igarashi, R. & Shirakawa, M. Machine-Learning Optimization of Multiple Measurement Parameters Nonlinearly Affecting the Signal Quality. ACS Meas. Sci. Au 1, 20–26 (2021).

36. Dickens, M. M., Bornhop, D. J. & Mitra, S. Removal of optical fiber interference in color micro-endoscopic images. Proc. 11th IEEE Symp. Comput.-Based Méd. Syst. (Cat 98CB36237) 246–251 (1998) doi:10.1109/cbms.1998.701364.

37. Kiyatkin, E. A. Brain temperature and its role in physiology and pathophysiology: Lessons from 20 years of thermorecording. Temperature 6, 271–333 (2019).

38. Higinio-Rodríguez, F., Rivera-Villaseñor, A., Calero-Vargas, I. & López-Hidalgo, M. From nociception to pain perception, possible implications of astrocytes. Front. Cell. Neurosci. 16, 972827 (2022).

39. Tegafaw, T. et al. Production, surface modification, physicochemical properties, biocompatibility, and bioimaging applications of nanodiamonds. RSC Adv. 13, 32381–32397 (2023).

40. Hollidge, B. S. et al. Kinetics and durability of transgene expression after intrastriatal injection of AAV9 vectors. Front. Neurol. 13, 1051559 (2022).

41. Orth, A., Ploschner, M., Wilson, E. R., Maksymov, I. S. & Gibson, B. C. Optical fiber bundles: Ultra-slim light field imaging probes. Science advances 5, (2019).

42. Chen, S., Liu, X. & McHugh, T. Near-infrared deep brain stimulation via upconversion nanoparticle-mediated optogenetics. Opt. Biopsy XVII: Towar. Real-Time Spectrosc. Imaging Diagn. 10873, 108730W–108730W–6 (2019).

43. Takahashi, M. et al. Investigating size and surface modification to optimise the delivery of nanodiamonds to brain glial cells. *Discov*. Nano 20, 143 (2025).

44. Chuma, S. et al. Implication of thermal signaling in neuronal differentiation revealed by manipulation and measurement of intracellular temperature. Nat. Commun. 15, 3473 (2024).

45. Hsieh, F.-J. et al. Bioorthogonal Fluorescent Nanodiamonds for Continuous Long-Term Imaging and Tracking of Membrane Proteins. ACS Appl. Mater. Interfaces 11, 19774–19781 (2019).

46. Wang, T. et al. Three-photon imaging of mouse brain structure and function through the intact skull. Nat. Methods 15, 789–792 (2018).

47. Rondin, L. et al. Magnetometry with nitrogen-vacancy defects in diamond. Rep. Prog. Phys. 77, 056503 (2014).

48. Sotoma, S. et al. Enrichment of ODMR-active nitrogen-vacancy centres in five-nanometre-sized detonation-synthesized nanodiamonds: Nanoprobes for temperature, angle and position. Sci. Rep. 8, 5463 (2018).

49. Chen, X.-D. et al. Temperature dependent energy level shifts of nitrogen-vacancy centers in diamond. Appl. Phys. Lett. 99, 161903 (2011).

